# Prefrontal activity sharpens spatial sensitivity of extrastriate neurons

**DOI:** 10.1101/2023.10.25.564095

**Authors:** M. Isabel Vanegas, Amir Akbarian, Kelsey L. Clark, William H. Nesse, Behrad Noudoost

**Affiliations:** Department of Ophthalmology and Visual Sciences, University of Utah, Salt Lake City, UT 84132, USA; Department of Mathematics, University of Utah, Salt Lake City, UT 84132, USA

**Keywords:** vision, feedback, primate, receptive field, FEF, V4, spatial sensitivity, inactivation, cortex, visual processing

## Abstract

Prefrontal cortex is known to exert its control over representation of visual signals in extrastriate areas such as V4. Frontal Eye Field (FEF) is suggested to be the proxy for the prefrontal control of visual signals. However, it is not known which aspects of sensory representation within extrastriate areas are under the influence of FEF activity. We employed a causal manipulation to examine how FEF activity contributes to spatial sensitivity of extrastriate neurons. Finding FEF and V4 areas with overlapping response field (RF) in two macaque monkeys, we recorded V4 responses before and after inactivation of the overlapping FEF. We assessed spatial sensitivity of V4 neurons in terms of their response gain, RF spread, coding capacity, and spatial discriminability. Unexpectedly, we found that in the absence of FEF activity, spontaneous and visually-evoked activity of V4 neurons both increase and their RFs enlarge. However, assessing the spatial sensitivity within V4, we found that these changes were associated with a reduction in the ability of V4 neurons to represent spatial information: After FEF inactivation, V4 neurons showed a reduced response gain and a decrease in their spatial discriminability and coding capacity. These results show the necessity of FEF activity for shaping spatial responses of extrastriate neurons and indicates the importance of FEF inputs in sharpening the sensitivity of V4 responses.

## Introduction

Frontal Eye Field (FEF) is believed to serve as a proxy for prefrontal cortex’s communication with posterior visual areas (Squire et al 2013). FEF sends direct projections to multiple visual areas, including extrastriate area V4 (Anderson et al 2011, Felleman & Van Essen 1991, Markov et al 2014, Schall et al 1995, Ungerleider et al 2008). FEF inactivation is associated with a reduction in an animal’s ability to detect sensory information in spatial attention tasks (Gregoriou et al 2014, Monosov & Thompson 2009, Thompson & Bichot 2005, Wardak et al 2012, Wardak et al 2006). Indeed, it has been shown that the strength of visual signals within the visual areas depends on the level of FEF activity (Armstrong et al 2006, Ekstrom et al 2009, Moore & Armstrong 2003, Noudoost & Moore 2011a, Ruff et al 2006, Schafer & Moore 2011, Silvanto et al 2006, Taylor et al 2007, Veniero et al 2021). Therefore, it has been suggested that the FEF’s role in covert attention (Armstrong et al 2009, Gregoriou et al 2009, Hamker & Zirnsak 2006, Moore & Fallah 2001, Zanto et al 2011), and overt attention (Bruce et al 1985, Robinson & Fuchs 1969, Schall 1991), relies on its ability to modulate the strength of visual signals in extrastriate areas (Buffalo et al 2010, Clark et al 2015, Moore et al 1998, Moran & Desimone 1985, Steinmetz & Moore 2014, Vinck et al 2013). In spite of the evidence for the role of FEF in gating feature-dependent signals in V4 and other visual areas, a direct measurement of how FEF contributes to the processing of spatial information within visual areas has never been conducted. FEF inactivation is associated with a reduction in the accuracy and precision of saccadic eye movements (Acker et al 2016, Dias et al 1995, Dias & Segraves 1999). Determining whether and how FEF shapes the spatial response profile of visual areas is important for our understanding of the circuitry involved in deployment of spatial attention. The goal of this study is to determine the degree to which the spatial profile of V4 receptive fields (RFs) depends on FEF activity.

Several considerations led us to test whether FEF contributes to shaping V4 neurons’ RF profile, even in the absence of an explicit attention or eye movement task. Inactivating FEF even only at the visual stimulus location during an oculomotor task increases scatter in saccade landing points, suggesting that FEF activity makes an important contribution to localizing visual targets (Acker et al 2016, Dias et al 1995, Dias & Segraves 1999). Earlier in the visual hierarchy, feedback from V2 to V1 helps shape RFs (Nurminen et al 2018); possibly feedback serves a similar role throughout the visual hierarchy. The fast visual latency of some FEF neurons (Pouget et al 2005, Schall 1991, Schmolesky et al 1998) suggests that their activity could contribute to initial visual responses in V4. Complicating this hypothesis, anatomical characterizations of the projections from FEF to V4 are not entirely consistent with either feedforward or feedback type projections (Anderson et al 2011, Ungerleider et al 2008). The strong reciprocal connections between FEF and V4 has placed both areas at a similar level in anatomically constructed hierarchies (Markov et al 2013). We specifically studied the RF profile of V4 neurons after FEF inactivation to understand the contribution of FEF-V4 reciprocal connections in shaping basic V4 response characteristics. In order to do that, we mapped the spatial response profile of V4 neurons before and after pharmacological inactivation of a retinotopically overlapping FEF site. We used Gaussian fits to quantify changes in the V4 neuron’s RFs. FEF inactivation increased both baseline and visually-evoked V4 activity, and increased the size of the RF. The net result of these changes was a reduction in the ability to localize visual stimuli based on the responses of V4 neurons. These results suggest that FEF activity is important for sensory processing within visual areas even in the absence of explicit cognitive or oculomotor demands.

## Results

In order to determine the role of FEF in shaping V4 responses, RFs of V4 neurons were measured using a grid of visual probe stimuli (Fig. 1a), before and after FEF inactivation. Visual probes were presented in a grid around the estimated RF location during fixation. Visual responses from 30 to 150ms following probe presentation were used for subsequent analyses (See methods for details). The RF of an example V4 neuron is shown in figure 1b. A neuron’s RF was characterized by its baseline, response amplitude, RF center (x_0_,y_0_) and RF spread (σ, see methods for details) using a 2-dimensional (2D) Gaussian fit. The Gaussian fit for the sample neuron in figure 1b is illustrated in figure 1c. In order to assess the influence of FEF on V4 RFs, we identified V4 neurons with RFs overlapping the FEF RFs, during a simultaneous recording (Fig. 1d). FEF RFs were estimated based on the endpoints of microstimulation-evoked saccades (Fig. 1e). Pharmacologically inactivating FEF, using GABA-a agonist muscimol, while recording V4 neurons allowed us to test the FEF’s contribution to shaping V4 neuronal responses. We employed a custom-made injectrode system for pharmacological inactivation (Noudoost & Moore 2011b, Vanegas et al 2019). The efficacy and spatial spread of FEF inactivation were assessed using a memory-guided saccade (MGS) task, which has previously been used to map the extent of FEF inactivation (Dias et al 1995, Dias & Segraves 1999). In the MGS task, the animal must remember the location of a visual cue throughout a delay period, and then saccade to the remembered location to receive a reward. Consistent with previous studies, following FEF inactivation, performance on the MGS task was disrupted in a spatially specific manner (Fig. 1f). The spatial extent of impaired performance increased over time following the infusion, and included multiple target locations within the visual hemifield contralateral the infusion-site RFs. Saccade endpoints were also more scattered for targets presented at the FEF RF (Fig. S2). These behavioral measures provided confirmation of FEF inactivation at the retinotopic site corresponding to the RFs of recorded V4 neurons. We recorded from 153 V4 units using single and array electrodes over 33 inactivation sessions (98 units, 24 sessions monkey 1; 55 units, 9 sessions monkey 2). Data analysis was restricted to the 111 units for which the 2D Gaussian provided a good fit (R^2^ goodness of fit = 0.76 ± 0.15; see Methods and Fig. S1a).

**Figure 1.**
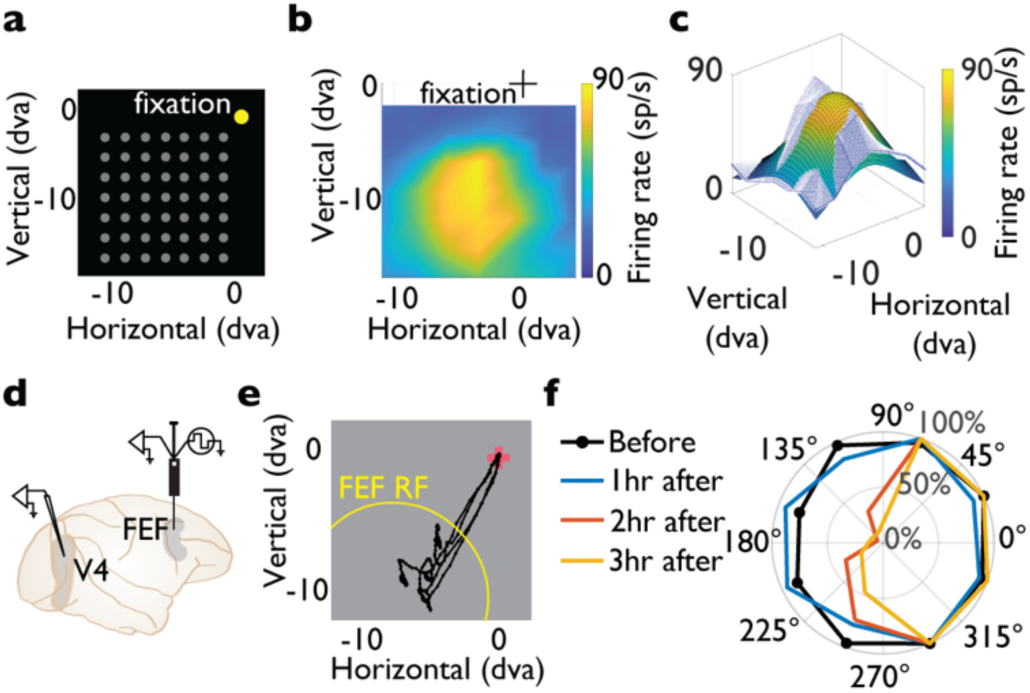
V4 RF mapping before and after FEF inactivation. **a**- Fixation task for V4 RF mapping. While the monkey fixated, brief white probes appeared pseudorandomly in a grid of possible locations. In this example, 49 possible locations are shown in gray. **b**- Example V4 neuron’s RF (before FEF inactivation). Color indicates average visual response across space. **c**- 3D plot of example V4 RF (white surface) and 2D Gaussian RF fit (color surface). **d**- V4 recordings were performed with a linear array or single electrode, before and after drug infusion via microinjectrode in FEF. **e**- Example eye position traces of microstimulation-evoked saccades, used to estimate the FEF RF. Traces are aligned to starting eye position. **f**- Example of animal’s performance on the MGS task at eight target locations, before and after FEF muscimol infusion. Colors indicate time relative to infusion (black shows before infusion, other colors 1-4hr after infusion). Before FEF infusion, performance is high at all locations. After muscimol infusion, performance drops for targets in the contralateral visual hemifield.

Figure 2 shows an example session during which V4 RFs were assessed before and after FEF inactivation. Figure 2a shows traces of eye position during the MGS. The eye position traces from the MGS task are overlaid on the V4 RF estimated based on visual responses during fixation. Following FEF inactivation, endpoints of saccades toward the overlapping FEF-V4 RF were more scattered (saccade scatter_before_=1.99±0.06 dva, saccade scatter_after_=3.29±0.07 dva, p=8.7×10^-36^; black traces in figure 2a). Saccades into the opposite hemifield were unaffected by the FEF inactivation (saccade scatter_before_=1.27±0.08 dva, saccade scatter_after_=1.39±0.03 dva, p=0.21; green traces in figure 2a). Figure 2b shows raster plots and firing rate for the V4 neuron in response to a probe presented at its RF peak, before and after FEF inactivation. Both the spontaneous activity of the neuron (prior to stimulus presentation) and the visually-evoked activity were greater following FEF inactivation (spontaneous activity_before_=33.73±10.65 sp/s, spontaneous activity_after_=74.87±8.22 sp/s, p=1.1×10^-3^; visual activity_before_=114.20±11.69 sp/s, visual activity_after_=178.69±5.16 sp/s, p=2.9×10^-6^). Thus, for this sample neuron, we found that FEF inactivation increases V4 activity in both the presence and absence of visual stimuli. Figure 2c shows the RF of the same neuron before and after FEF inactivation, overlaid with the corresponding contour profiles based on 2D Gaussian fits (R^2^_before_=0.87, R^2^_after_=0.93). We found that after FEF inactivation, V4 RF expanded: the RF spread parameter of the 2D Gaussian changed from 3.16 dva before inactivation to 3.79 dva, after. Thus, the response of this example V4 neuron became less spatially precise following FEF inactivation. Figure 2d shows a horizontal cross-section of the V4 RF at its peak and the corresponding Gaussian fit. Using the Gaussian fit parameters, we found that not only the peak value at the RF center was increased, the baseline value for probes farther away from the RF center was also increased. Baseline value changed from 35.49 sp/s to 58.77 sp/s after inactivation and peak value at the RF center changed from 101.29 sp/s to 174.58 sp/s. The center of the RF moved very little following FEF inactivation, from RF coordinates of [-4.31, −7.39] dva before to [-3.22, −7.81] dva after inactivation, resulting in a total shift of 1.16 dva.

**Figure 2.**
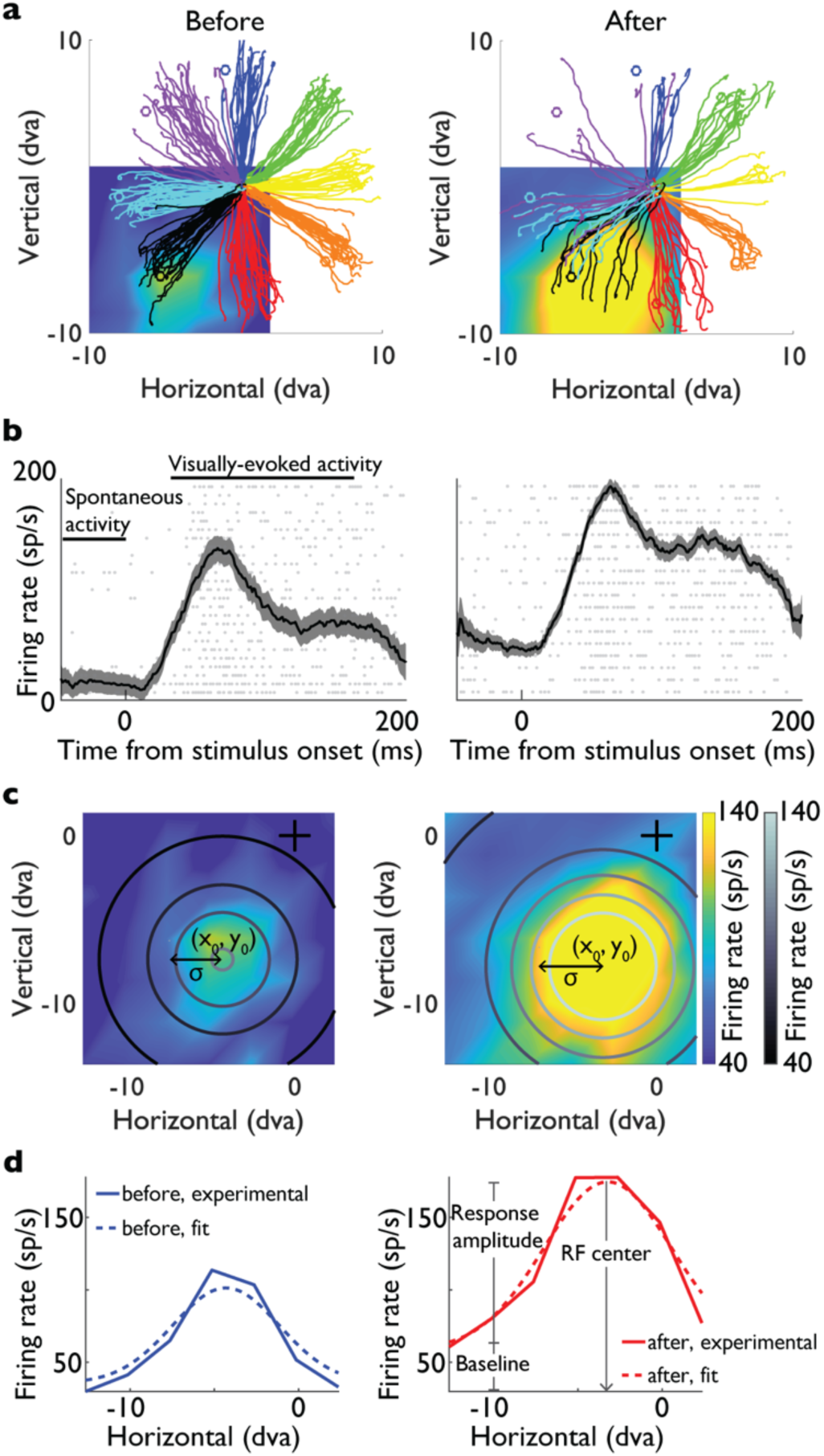
Responses and 2D Gaussian fits of an example V4 neuron before and after FEF inactivation. **a**- Eye movement traces for memory guided saccades before (left) and after (right) FEF inactivation. Colors correspond to the eight target locations. Heat map in the background shows the example V4 RF, near the endpoints of the black traces. **b**- Visually-evoked activity at the peak RF location, before (left) and after FEF inactivation (right). Raster plots: each row represents one trial, each gray dot represents an action potential. Overlaid are firing rates (peri-stimulus time histogram, or PSTH traces, black) in response to probe at the location evoking the maximum V4 response, before (left) and after FEF inactivation (right). Plot shows mean ± SE across trials. Horizontal black lines indicate the time range used to calculate the spontaneous and visually-evoked activity. **c**- V4 RF before (left) and after FEF inactivation (right) overlaid with contour profiles based on 2D Gaussian fits. Color indicates visually-evoked activity across horizontal and vertical probe locations. σ indicates the spread of the Gaussian, or RF spread. The center of the Gaussian, or RF center, is at (x_0_,y_0_). Black cross indicates fixation point in all RF plots. **d**- Cross-sectional views of the experimental (solid) and Gaussian (dashed) V4 RF, before (left, blue) and after FEF inactivation (right, red). Cross-section shows responses at different horizontal probe positions for −6.1 dva vertical (chosen to include peak overall response).

These changes in V4 activity and RF properties following FEF inactivation were replicated at the level of the population (n=111 V4 neurons). The normalized activity of V4 population are shown in figures 3a. Both the spontaneous activity and the visually-evoked activity in V4 increased following FEF inactivation (spontaneous activity_before_=0.11±0.01, spontaneous activity_after_=0.22±0.01, p=2.8×10^-11^; visual activity_before_=0.39±0.02, visual activity_after-_=0.51±0.02, p=3.5×10^-7^). Figure 3b shows the average spatial profile of the 2D Gaussian fits based on the normalized response of V4 population. Following FEF inactivation, the baseline and RF spread increased (baseline_before_=0.02, baseline_after_=0.25, RF spread_before_=2.71 dva, RF spread_after_=2.86 dva). However, the response amplitude and RF center were unchanged (response amplitude_before_=0.66, response amplitude_after_=0.67, RF center (x,y)_before_=(−0.97,-3.05), RF center (x,y)_after_=(−0.94,-3.09). The same pattern was observed across the Gaussian fits based on the raw response of V4 neurons as well. FEF inactivation increased the baseline activity (Fig. 3c, baseline_before_=27.19±3.26 sp/s, baseline_after_=37.64±3.60 sp/s, p=1.3×10^-11^). RF spread also increased following FEF inactivation (Fig. 3d, RF spread_before_=2.71±0.09 dva, RF spread_after_=2.86±0.08 dva, p=9×10^-5^). However, FEF inactivation did not alter response amplitude of V4 neurons (Fig. 3e, response amplitude_before_=38.37±4.74 sp/s, response amplitude_after_=40.26±4.66 sp/s, p=0.35). RF centers also remained unchanged following inactivation (Fig. S1b; RF center_x, before_=-0.97±0.54 dva, RF center_x, after_=-0.94±0.55 dva, p=0.55, and RF center_y, before_=-3.05±0.25 dva, RF center_y, after_=-3.09±0.25 dva, p=0.61).

**Figure 3.**
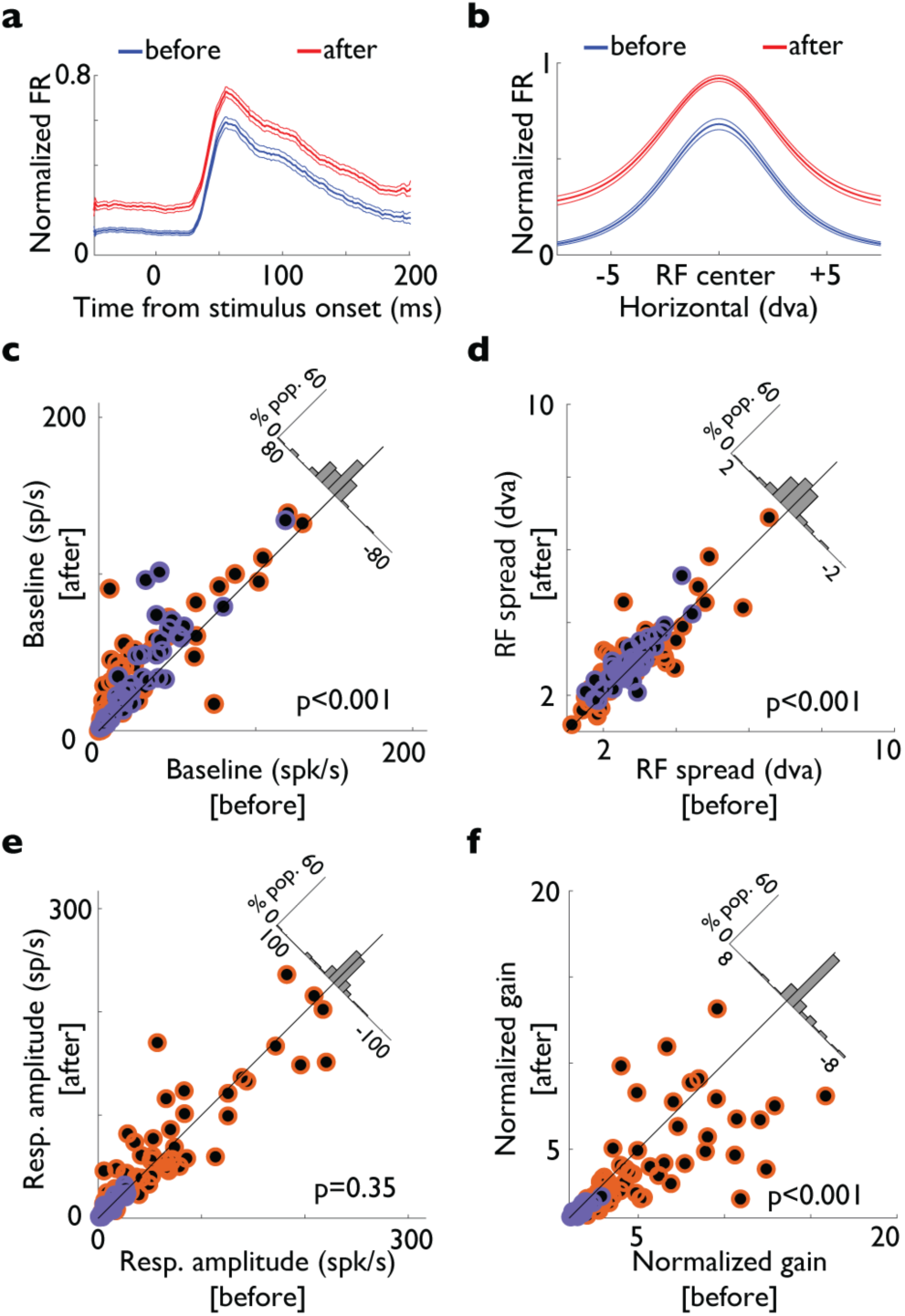
Changes in RF properties of the V4 population following FEF inactivation. **a**- Normalized population visually-evoked activity before (blue) and after FEF inactivation (red). Response is to best probe location, plotted as mean ± SE across 111 neurons. **b**- Population mean Gaussian fit before (blue) and after FEF inactivation (red). Response amplitude and baseline are shown for the after fit. **c-f**: Parameters extracted from individual 2D Gaussian fits before vs. after FEF inactivation, for 69 neurons in monkey 1 (orange) and 42 neurons in monkey 2 (purple): **c**- Baseline, **d**- RF spread (σ), and **e**- Response amplitude, **f**- Normalized gain. Histograms in the upper right show the difference (after-before) of each parameter value. Two units with baseline values >140sp/s are not shown, however they are included in the statistical tests.

Several lines of evidence indicate the role of FEF in boosting visual signals in posterior visual areas (Armstrong et al 2006, Armstrong & Moore 2007, Noudoost & Moore 2011a). The finding that FEF inactivation enhances V4 signals in terms of spontaneous activity, visual-evoked activity is counterintuitive. On the other hand, from a signal detection perspective, the finding that after FEF inactivation, baseline activity is enhanced but response amplitude is not, raises the possibility that FEF inactivation mostly enhances the noise rather than affecting the visual signal. In other words, FEF inactivation might alter the gain of visual signals in terms of signal to noise ratio at the level of V4. In order to assess the capacity of visual neurons to process sensory information, we quantified the normalized gain as the ratio: (baseline+response amplitude) / baseline. Higher normalized gain values correspond to a higher signal-to-noise ratio. We found that FEF inactivation strongly reduced the normalized gain of V4 responses (Fig. 3f, normalized gain_before_=3.70±0.29, normalized gain_after_=2.96±0.23, p=3.1×10^-9^). See figure S3 for non-uniform Gaussian parameter fits. The finding that after FEF inactivation V4 neurons respond more in the absence of robust visual stimuli (enhanced baseline and spontaneous activity), suggest that FEF might play its role in boosting sensory information by sharpening the response to visual signals while suppressing the response to sub-optimal ones.

We used our Gaussian fits as an estimate of the firing rate, to compute a theoretic information measure of discrimination information (DI), also known as Kullback-Leibler divergence, or relative entropy. Assuming a Poisson process for the neuron’s spike generation, DI was calculated as:

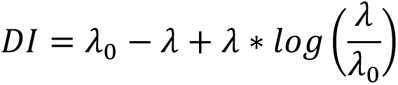

where λ_0_ is the baseline firing rate, and λ is baseline + response amplitude based on individual Gaussian parameter fits.

Figure 4a shows the DI before and after FEF inactivation, for stimuli presented at the RF center. V4 population showed a significant drop in their ability to detect RF stimuli (DI_before_=27.39±4.08 bits, DI_after_=23.95±3.77 bits, p=2.3×10^-3^). Figure 4b shows population average change in DI (after-before), as a function of stimulus distance from the RF center. DI was significantly reduced following FEF inactivation for locations within 1.9 dva of the RF center (p<0.05). Figure 4c shows the change in DI due to FEF inactivation based on the average 2D Gaussian fit of V4 neurons. Near the RF center there was a decrease of up to 15% in the DI values following FEF inactivation. Locations far from the RF center showed no change in DI. These results show that the changes in baseline response of V4 neurons after FEF inactivation can potentially result in a reduced capacity of these neurons to detect stimuli appearing around their RF, based on an information theoretic measure.

**Figure 4.**
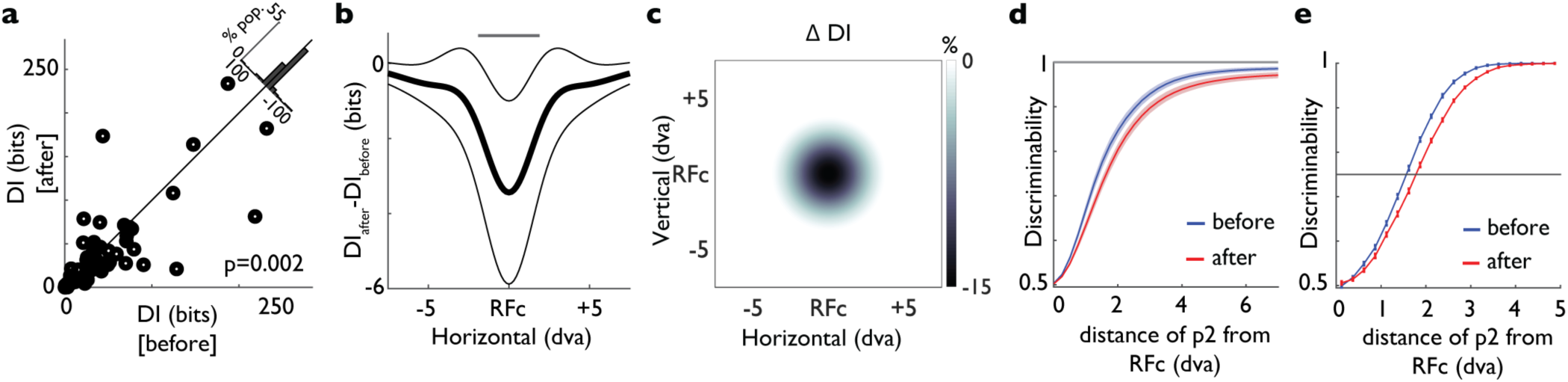
Spatial precision of V4 responses decreases following FEF inactivation. **a-** DI before and after FEF inactivation, based on the fitted Gaussian for each neuron. Histogram (upper right) shows the distribution of differences (after - before). **b-** Change in DI (after-before) as a function of distance from RF center (mean ± SE across the population of individual Gaussian fits). Gray bar on top shows the range where change is significant (p<0.05). **c-** Percent change in DI across space relative to the RF center, based on the average Gaussian parameters across the population. **d-** Two-point discriminability as a function of distance between stimuli, before (blue) and after FEF inactivation (red), plotted as mean ± SE across the population of individual Gaussian fits. Discriminability was measured using ROC analysis on firing rates generated from the Gaussian fits, with one probe at the RF center and the second probe a variable distance away. **e-** Same as d, for population average Gaussian fit.

We also employed a receiver operating characteristics (ROC) method to assess the V4’s capacity to differentiate between pairs of stimuli one appearing at the RF center and appearing at different locations (two-point discriminability; see Methods and Fig. S4b for details). Figure 4d shows the average two-point discriminability, computed based on the V4 responses. Following FEF inactivation, average discriminability decreased for all stimuli farther than 0.01 dva from the RF center (p<0.002, n=111 neurons). Using the Gaussian fit we also observed a similar phenomenon (Fig. 4e). Stimuli farther from the RF center were less discriminable after FEF inactivation than before. The distance at which half of discriminability performance was reached (75% discriminability) increased from 1.57 dva to 1.78 dva after FEF inactivation (95% CI [1.56,1.58] dva before, 95% CI [1.75, 1.81] dva after). Moreover, the slope of the logistic function also decreased from 2.24 to 1.96 after FEF inactivation (95% CI [2.20,2.28] before, 95% CI [1.88,2.04] after). Altogether, these results indicate that after FEF inactivation, the ability of V4 neurons to discriminate the location of two stimuli around their RF is reduced.

## Discussion

We assessed V4 RF characteristics before and after pharmacological inactivation of retinotopically corresponding FEF sites (Fig. 1). Following FEF inactivation, both baseline and peak firing rates in V4 increased, and RF size expanded (Fig. 2&3). The combination of increased baseline firing rate and peak firing rate resulted in a decrease in the overall gain of the visual response. Consequently, following FEF inactivation, we found a decrease in the amount of information in V4 about the stimulus location (Fig. 4). Altogether, these results indicate that FEF activity plays an important role in shaping extrastriate visual responses, even during a passive fixation task, without any explicit cognitive demands. Our finding that FEF inactivation reduces visual information in extrastriate cortex is broadly consistent with a large body of prior literature showing that increases in FEF activity increase visual information in these areas. Microstimulation of FEF increases feature discrimination in V4 (Armstrong & Moore 2007, Moore & Armstrong 2003), as does activating FEF neurons with dopaminergic manipulations (Noudoost & Moore 2011a). Similarly, inactivating FEF reduced the increase in visual discrimination that usually accompanies saccades (Moore et al 1998, Noudoost et al 2014). Moreover, during working memory, while V4 neurons are receiving input from FEF, their representation of the location of visual stimuli is enhanced (Merrikhi et al 2017). In line with these previous results, here we showed that in the absence of FEF input, information about the location of visual stimuli is reduced.

While our results confirm the contribution of FEF to the gating of visual signals, the observed effects of inactivating FEF are not merely opposite those of activating it. Stimulating FEF increases information in visual cortex through a preferential increase in firing rate for preferred stimuli (Armstrong & Moore 2007, Moore & Armstrong 2003, Noudoost & Moore 2011a); in our findings, inactivating FEF also leads to an increase in extrastriate firing rates, but in a nonspecific way which reduces information. Therefore, based on our results, rather than a reduction in gain, FEF inactivation seems to release V4 from a global inhibition, resulting in nonspecific response increases. In other words, FEF activity sharpens the spatial selectivity of V4 neurons.

The direct projections from FEF to V4 are an obvious candidate for mediating the influence of FEF activity on V4 visual responses; here we briefly review what is known about these projections. Projections from FEF to V4 terminate across all layers, and arise predominantly from the superficial layers of FEF, and are therefore described as being of an ‘intermediate’ rather than purely feedforward or feedback type (Anderson et al 2011, Ungerleider et al 2008). V4- projecting FEF neurons are characterized by delay activity during a working memory task (Merrikhi et al 2017) (some also have visual activity) and express D1R receptors (Mueller et al 2020)– consistent with the ability of FEF D1R manipulation to alter both delay activity within FEF (Williams & Goldman-Rakic 1995) and visual selectivity in V4 (Noudoost & Moore 2011a).

Synapses from these FEF projections are primarily (98%) onto putatively excitatory neurons within V4 (Anderson et al 2011). Based on these anatomical facts, there are at least three potential mechanisms which could drive the observed changes in V4 responses. In both of the first two scenarios, despite mainly targeting pyramidal neurons (Anderson et al 2011), our observed changes in V4 could be accounted for by FEF projections that preferentially activate inhibitory neurons. This idea is also supported by the finding that inhibitory neurons in V4 show larger firing rate changes than excitatory neurons during attention (Mitchell et al 2007) and working memory (Nesse et al 2021). Such selective activation of inhibitory neurons has also been shown to sharpen feature selectivity of excitatory neurons in V1 (Lee et al 2012). Loss of such inhibitory activity could explain the expanded excitatory RFs following FEF inactivation, as inhibitory activity is responsible for creating the inhibitory surround (Adesnik et al 2012); elsewhere in the visual hierarchy, removing feedback from V2 to V1 similarly increases responses in the RF surround (Nassi et al 2013, Nurminen et al 2018). This activation of inhibitory V4 neurons from FEF input could either be mediated 1) via a strong influence of the few FEF synapses onto inhibitory neurons, or 2) by a small intermediate excitatory V4 population which in turn recruit local inhibitory networks to alter V4 visual responses. However, a third possibility, that the FEF inactivation causes indirect compensatory effects in other connected areas, cannot be ruled out; for example, the lateral intraparietal area (LIP) might undergo a compensatory increase in activity following FEF inactivation (analogous to (Balan et al 2019)), and that increased activity in the LIP projections to V4 (Markov et al 2014, Ungerleider et al 2008) might then drive the nonspecific increase in V4 activity. Therefore, although our results show the contribution of FEF to sharpening spatial tuning within V4, further work will be needed to determine the specific circuitry involved in prefrontal control of visual signals.

## Materials and methods

### General and surgical procedures

Two male rhesus monkeys (Macaca mulatta, 12 and 16Kg) were used in these experiments. All experimental procedures were in compliance with the National Institutes of Health Guide for the Care and Use of Laboratory Animals, the Society for Neuroscience Guidelines and Policies, and the University of Utah Institutional Animal Care and Use Committee. Each of the animals underwent surgery to implant a head post and two cylindrical recording chambers (20mm diameter). Stereotactic surgery coordinates were (AP 25+(2), ML 15 (±0)) and (AP −5(−1),ML 20+(2)) of the right hemisphere for monkey 1, and (AP 30+(1-2), ML 15-(1-2)) and (AP −5-(1-2), ML 20+(1-2)) of the left hemisphere for monkey 2. Two craniotomies were performed on each animal, allowing access to the FEF area on the anterior bank of the arcuate sulcus, and dorsal extrastriate visual area V4 on the prelunate gyrus. All surgeries were conducted under sterile techniques and general anesthesia (isoflurane), with analgesics during the post-surgery recovery period.

### Behavioral tasks

All visual tasks were programmed using the NIMH Monkeylogic toolbox (ML2) (Hwang et al 2019), on 64-bit Matlab (The MathWorks, Inc., Natick, MA). In all tasks, we monitored eye position with an infrared optical eye-tracking (EyeLink 1000, SR Research, Ottawa, Canada), in a distress-free setup in which a “hot” (infrared) mirror was used to reflect the monkey’s eye position to the camera. All tasks were projected on a VG248 ASUS LED monitor positioned 33cm in front of the animal, with a refresh rate of 144Hz and resolution of 1920 x 1080 pixels. See SI for details of FEF and V4 RF estimation, and MGS performance and scatter used to assess the efficacy of FEF inactivation.

#### Visual tasks for V4 RF mapping

On the day of the FEF inactivation experiment, we first estimated V4 RFs based on audible responses to oriented bars. Then we quantitatively measured V4 RFs using a series of stationary stimuli (probes) on a black background. Probes were pseudorandomly presented, based on a grid of possible locations, centered on the estimated V4 RF center based on prior V4 RF localization. The monkey had to maintain fixation (window ±1.5 dva monkey 1, ±2 dva monkey 2) throughout the full trial for data to be included.

For monkey 1, probes were white circles (1dva diameter), 100ms duration, 100ms between probes, chosen from 49 locations on a 7×7 grid (2.5 dva spacing) extending over 15×15 dva. In a given trial, we presented eight probes, with an inter-trial interval of 1000ms. The task required fixation on a central white circle (1dva diameter) throughout the visual stimulation period, in order to obtain a juice reward.

For monkey 2, probes were white squares (0.8×0.8dva), 7ms duration, 0ms between probes, pseudorandomly chosen from 81 locations on a 9×9 grid (4.3 dva spacing) extending over 25×25 dva. In this task, fixation was required for at least 1000ms on a central white circle (0.75 dva diameter). Upon disappearance of the fixation spot, a saccade was made towards a target placed on the left side of the screen. In order to avoid perisaccadic influence on the RF mapping, we only analyzed data from 1000ms to 500ms before the saccade onset.

### Neurophysiological recordings

Data acquisition was performed via Neuralynx and Blackrock systems. Spike waveforms were digitized and stored at 32 kHz for offline spike sorting and data analysis. Extracellular waveforms were digitized and manually sorted (Offline Sorter, Plexon). V4 recordings were made through the surgically implanted chamber. Electrodes were placed in the cortex using a hydraulic microdrive (Narishige, Japan). We recorded from V4 neurons using single tungsten microelectrodes of 200μm diameter, with epoxylite insulation (FHC, Bowdoin, ME), and linear 16-channel arrays with 150μm space between contacts (Plexon, Dallas, TX). We performed drug infusion using a custom-made microinjectrode, described in previous publications (Noudoost et al 2014, Vanegas et al 2019).

### Inactivation of the FEF

We pharmacologically inactivated FEF by infusing 0.5-1μL of the GABA-a agonist muscimol, as described in (Noudoost & Moore 2011b), and demonstrated more recently in (Noudoost et al 2014, Vanegas et al 2019). In the example session we injected a volume of 1μL. The concentration of muscimol was 5mg/ml (pH 6.5 to 7) (Noudoost et al 2014). We did not perform inactivation experiments on consecutive days. Details of MGS performance and scatter used to assess the efficacy of FEF inactivation are in the SI.

### Data analyses

All analyses were programmed on 64-bit Matlab 2018b (The MathWorks, Inc., Natick, MA). Unless otherwise specified, all p values were based on the Wilcoxon signed rank test (paired samples). A criterion of p<0.05 was established for significance. We performed 33 FEF inactivation sessions (24 in monkey 1, and 9 in monkey 2), in which a total of 153 V4 units (98 in monkey 1, 55 in monkey 2) were recorded.

#### V4 RF analyses

V4 RF maps were calculated from average firing rate responses to the visual probes, in the time window from 30-150ms after stimulus onset (Fig 2b, range within pink lines), in spikes per second (sp/s, Fig 2c). On average, the number of trials per stimulus location were: 20±1 and 20±2 for monkey 1, and 712±4 and 706±6 for monkey 2, before and after FEF inactivation, respectively.

For the location with the maximum response to the visual stimulus (RF peak), we computed the firing rate over time, or peri-stimulus histogram. Here, we applied a moving average smoothing filter with a window size of 20ms (Fig 2b, 3a).

For the population PSTH (Fig 3a) we normalized the responses of each neuron according to the following formula:

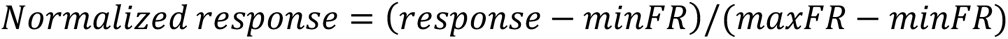

Where maximum and minimum firing rates were calculated across all times and conditions before and after FEF inactivation (in a 20ms bin, after smoothing).

#### 2D-Gaussian fit to V4 RF data

We performed a two-dimensional (2D) Gaussian fit of the V4 RF data using least squares optimization. The 2D Gaussian function is expressed as:

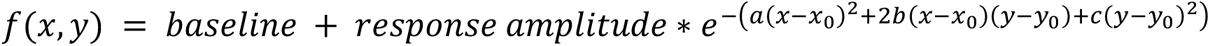

Where a, b and c account for the rotation of the 2D Gaussian by a clockwise angle θ:

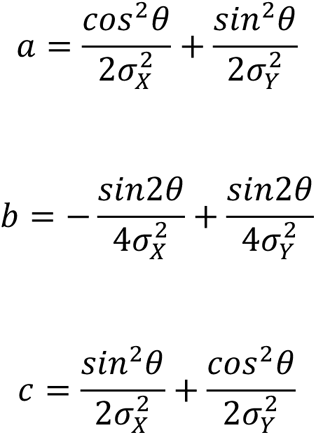

The coefficients *baseline* and *response amplitude* capture the baseline firing rate and amplitude of the RF, the coordinate (x_0_,y_0_) is the RF center, and σ_X_ and σ_Y_ are the RF spread in a two-dimensional space.

When the RF spread is uniform, σ_X_ = σ_Y_ = σ. The 2D Gaussian function can be simplified to:

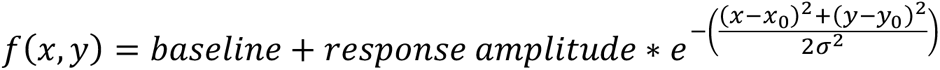

Initialization parameters for the center of the Gaussian, baseline, and response amplitude, were chosen from the center of the RF, minimum and maximum RF response, measured in experimental data for each neuron individually. Lower bounds on the coefficients to be fitted for the baseline, response amplitude, and RF spread, were set at zero. For the center of the Gaussian, lower bounds were the minimum coordinate points according to the extent of visual stimulation in dva in the session. Upper bounds were set at the maximum RF response for the baseline, and twice the maximum RF response for the response amplitude. For RF spread, the upper boundary was based on the extent of visual stimulation in dva (monkey 1), and we used a more constrained boundary of 10dva for monkey 2. For the center of the Gaussian, upper bounds were the maximum coordinate points according to the extent of visual stimulation in dva in the session. We also applied a 2D interpolation of the experimental RF (8 points between sample values) to smooth and enhance the RF resolution before applying the 2D Gaussian fit.

For the next step of our analysis (population before vs after FEF inactivation comparisons, spatial precision using discrimination information and ROC), we included, based on the individual 2D- Gaussian fit, 111 units for which the R^2^ goodness of fit was greater than 0.6 for monkey 1 and 0.4 for monkey 2 (Fig. S1a-b, 12 single units and 57 multi-unit recordings for monkey 1, and 42 single units for monkey 2).

The population average Gaussian fit (Fig 3b) was calculated by fitting a Gaussian to the normalized response of each neuron and then averaging the parameters.

Spontaneous and visually-evoked activity were calculated in a 20-ms window from 1 to 20ms, and in a 120-ms window from 30 to 150ms following the stimulus presentation, respectively.

#### Spatial precision of V4 responses

We used our Gaussian fits as an estimate of the firing rate, to compute a theoretic information measure of discrimination information (DI), also known as Kullback-Leibler divergence, or relative entropy. Assuming a Poisson process for the neuron’s spike generation, DI was calculated as:

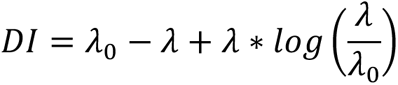

where λ_0_ is the baseline firing rate, and λ is baseline + response amplitude based on individual Gaussian parameter fits.

DI was computed individually for each neuron, at the RF center location, before vs. after FEF inactivation (Fig 4a), and as the average difference (DI_after_-DI_before_, Fig 4b). We also calculated the percent change in DI as a function of stimulus locations relative to V4 RF center. In this case, each (x,y) location estimate for λ was computed from the average Gaussian parameters, before and after FEF inactivation (Fig 4c). Change in DI was calculated as ΔDI = (DI_after_-DI_before_)/(max DI_before_-min DI_before_) %.

#### Discriminability with ROC analysis

The discriminability measure, *D*, was defined as the ability of an observer to dissociate between two probes, presented at a certain distance apart (*Δx*), based on the neuron’s predicted response (*λ*). The fitted Gaussian parameters were used to predict the neuron’s response to each of a pair of probes, one at the center of the RF (i.e., at *x*_%_, *y*_%_) and the other one at a distance of *Δx* (i.e., at *x*_%_ + *Δx*, *y*_%_). Based on the predicted firing rate for each probe location (calculated based on the fitted Gaussian function above), a distribution of neural responses was generated by adding a normally distributed random noise with a standard deviation of 10 percent of the predicted firing rate. The discriminability measure was then defined as the area under the curve (AUC) of the ROC between the predicted firing rate distributions for the two probe locations. For each distance between probes, 100 sample firing rates were generated to form the distributions and the AUC values were calculated for various distance values in 0.01 dva steps. Figure 4d uses Gaussian fits for individual neurons, and error is calculated over the population. Figure 4e uses the population average parameters for a Gaussian fit, and the error bars are over distances ±.5 dva from the point plotted.

#### Logistic function fit

We fit the discriminability curves from 4e with the following logistic function:

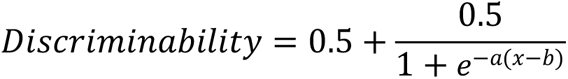

b is the value of the sigmoid’s midpoint (half performance)

1 is the curve’s maximum value

a is the logistic growth rate or steepness of the curve

### Statistical tests

All statistical comparisons used the Wilcoxon signed-rank test for paired comparisons and the Wilcoxon ranksum test for unpaired comparisons unless otherwise specified in the text.

## Data and code availability

All analyses were performed in Matlab using custom scripts. All data and code for analyses will be posted on GitHub [https://github.com/isabelvanegas] upon publication.

## Acknowledgements

We thank the veterinarian staff at the University of Utah, and Rochelle Moore for assistance with animal care. We thank Tyler S. Davis for assistance during surgical procedures. This work was supported by funding from the National Institutes of Health (NIH), grants R01EY026924, R01NS113073, and EY014800 to B.N., and an Unrestricted Grant from Research to Prevent Blindness, New York, NY, to the Department of Ophthalmology & Visual Sciences, University of Utah.

## Author contributions

M.I.V. and B.N. conceived the project. B.N. performed the surgical procedures. M.I.V. and A.A. performed experiments and collected electrophysiological data. M.I.V. and A.A. performed electrophysiology data analysis. M.I.V., A.A., W.H.N., and B.N. developed the methodology for mathematical modeling. M.I.V., K.L.C. and B.N. wrote the manuscript. All authors contributed to final edits of the manuscript.

## Supplemental information

### Prefrontal activity sharpens spatial sensitivity of extrastriate neurons

M. Isabel Vanegas, Amir Akbarian, Kelsey L. Clark, William H. Nesse, Behrad Noudoost

**Figure S1.**
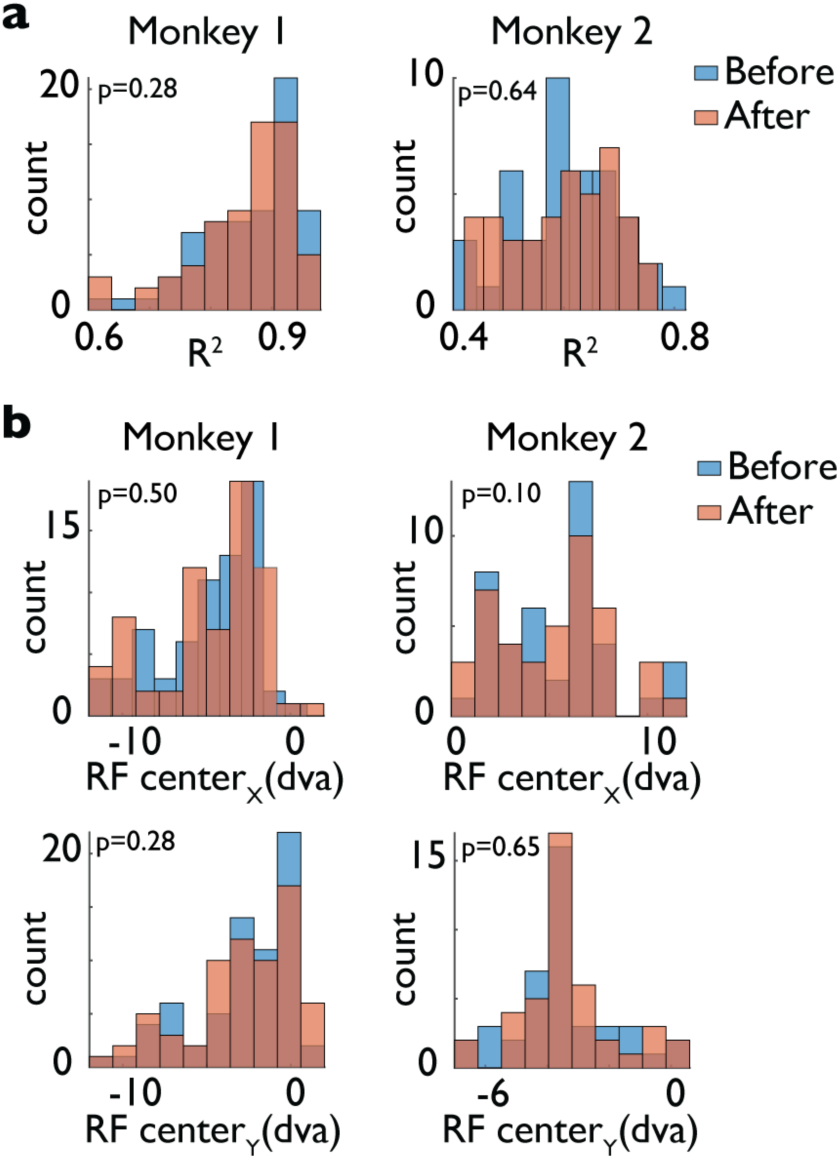
Consistency of RF fits and centers after FEF inactivation. **a.** Goodness of fit measure (R^2^) for the 2D Gaussian model in Monkey 1 (left) and Monkey 2 (right), before (blue) and after FEF inactivation (red). **b.** Center of V4 RF for all neurons, in the horizontal (top) and vertical dimensions (bottom), in Monkey 1 (left) and Monkey 2 (right), before (blue) and after FEF inactivation (red). In all plots, p values indicate comparisons of before vs. after. Monkey 1: R^2^_before_=0.87±0.01, R^2^_after_=0.87±0.01, p=0.28 Monkey 2: R^2^_before_=0.65±0.02, R^2^_after_=0.66±0.02, p=0.64 Monkey 1: RF center x_before_=-5.20±0.33, RF center x_after_=-5.17±0.36, p=0.50 Monkey 2: RF center x_before_=5.02±0.57, RF center x_after_=5.38±0.53, p=0.10 Monkey 1: RF center y_before_=-2.89±0.34, RF center y_after_=-2.98±0.35, p=0.28 Monkey 2: RF center y_before_=-3.49±0.26, RF center y_after_=-3.56±0.27, p=0.65

**Figure S2.**
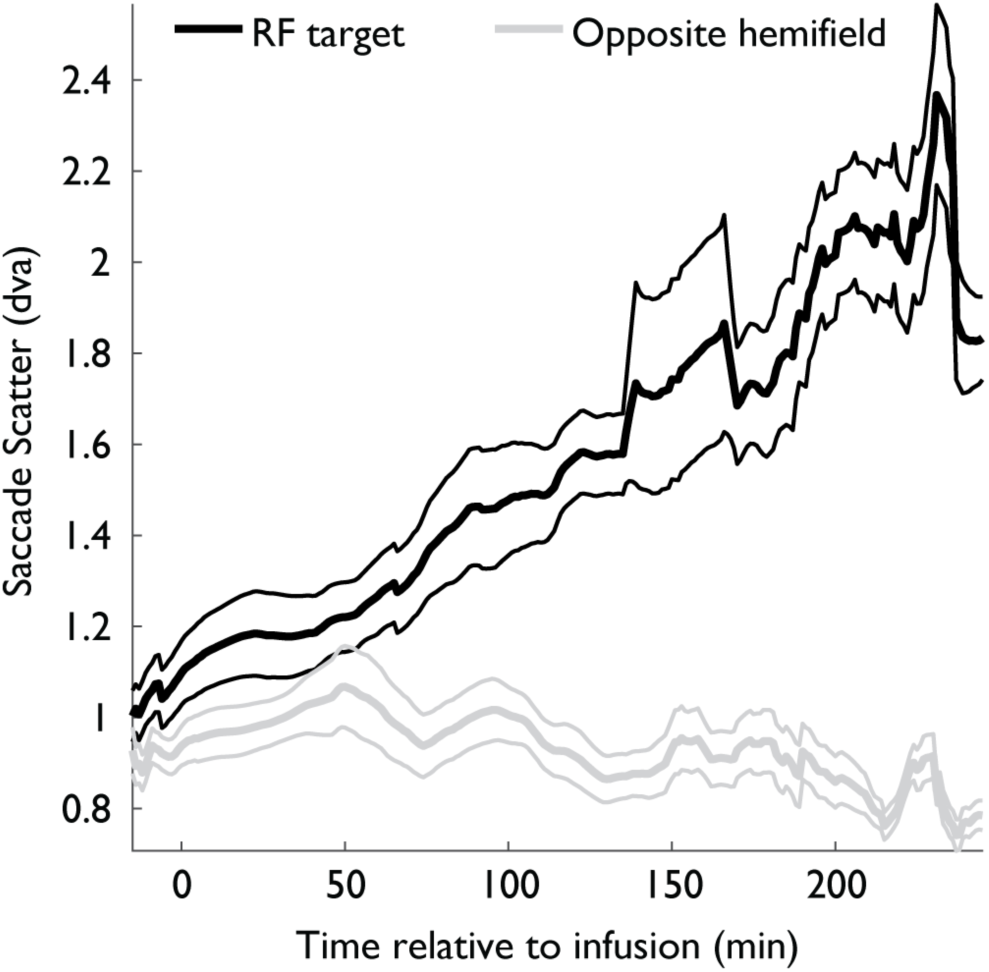
Eye position and saccades before and after FEF inactivation. Scatter of saccade endpoints in the MGS task, over time, relative to FEF inactivation. Data are shown for saccades with the memory (target) location in the RF (black) or in the opposite hemifield (gray). Scatter was measured for a moving window of 6 trials. Plot shows mean ± SE across.

**Figure S3.**
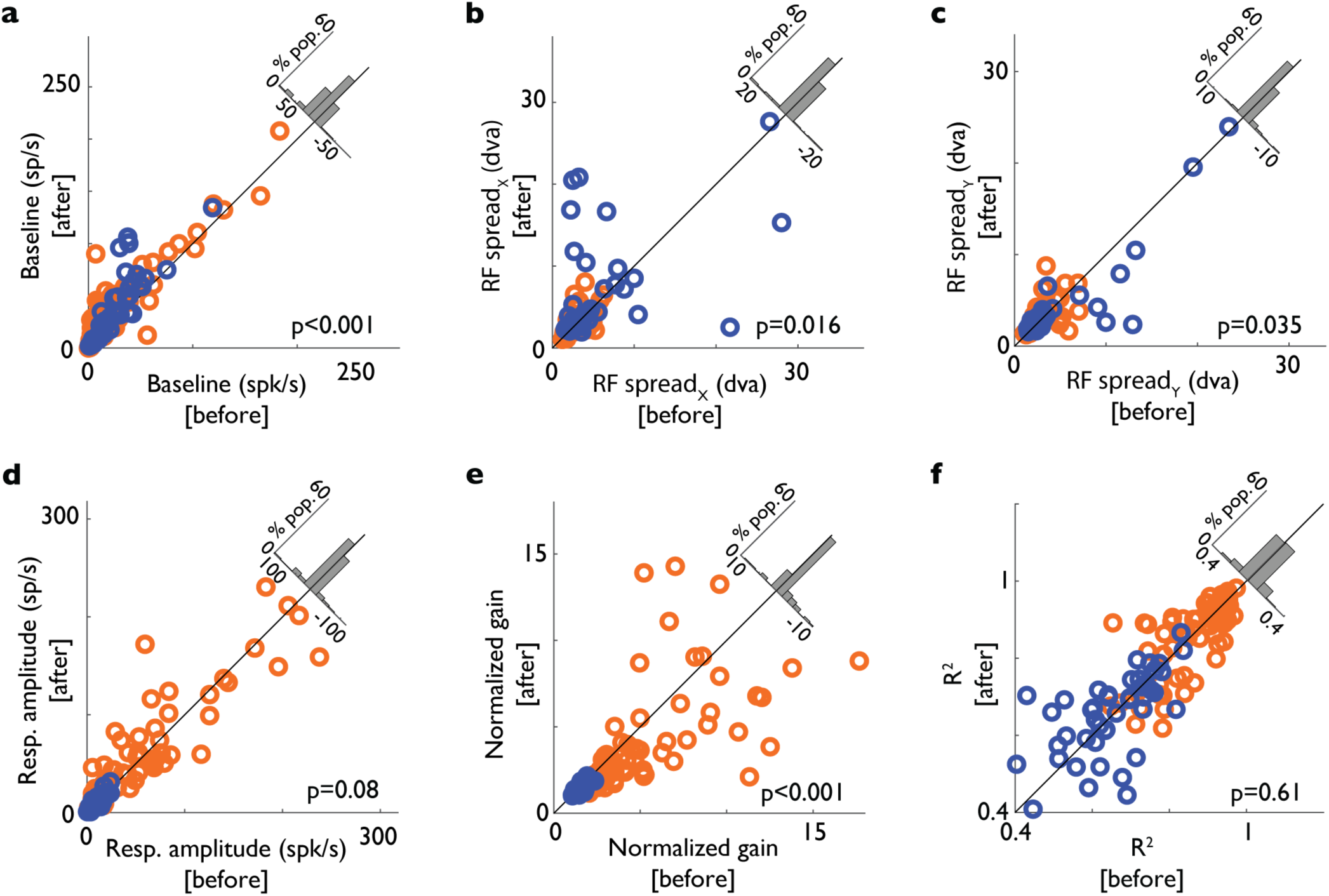
Parameters for non-uniform Gaussian before and after FEF inactivation. Each plot shows one parameter value before vs. after FEF inactivation for Monkey 1 (blue) and Monkey 2 (orange). Histograms in the upper right show change across neurons (after - before). Parameters were baseline (**a**), RF spread_X_ (**b**), RF spread_Y_ (**c**), response amplitude (**d**), normalized gain (**e**), along with R^2^ goodness of fit (**f**). p values compare before vs. after. Consistent with the results seen for the uniform 2D Gaussian fit, FEF inactivation increased baseline firing rate (baseline_before_=25.37±2.97 sp/s, baseline_after_=36.35±3.35 sp/s, p=2.2×10^-13^) and RF spread values (RF spread x_before_=3.74±0.35 dva, RF spread x_after_=4.18±0.37 dva, p=0.016; RF spread y_before_=3.43±0.28 dva, RF spread y_after_=3.34±0.25 dva, p=0.035), did not alter response amplitude (response amplitude_before_=35.08±4.43 sp/s, response amplitude_after_=37.15±4.30 sp/s, p=0.08), and resulted in a drop in normalized gain (normalized gain_before_=3.60±0.29, normalized gain_after_=2.95±0.24, p=4.6×10^-9^). R^2^_before_=0.79±0.01, R^2^_after_=0.79±0.01, p=0.61.

**Figure S4.**
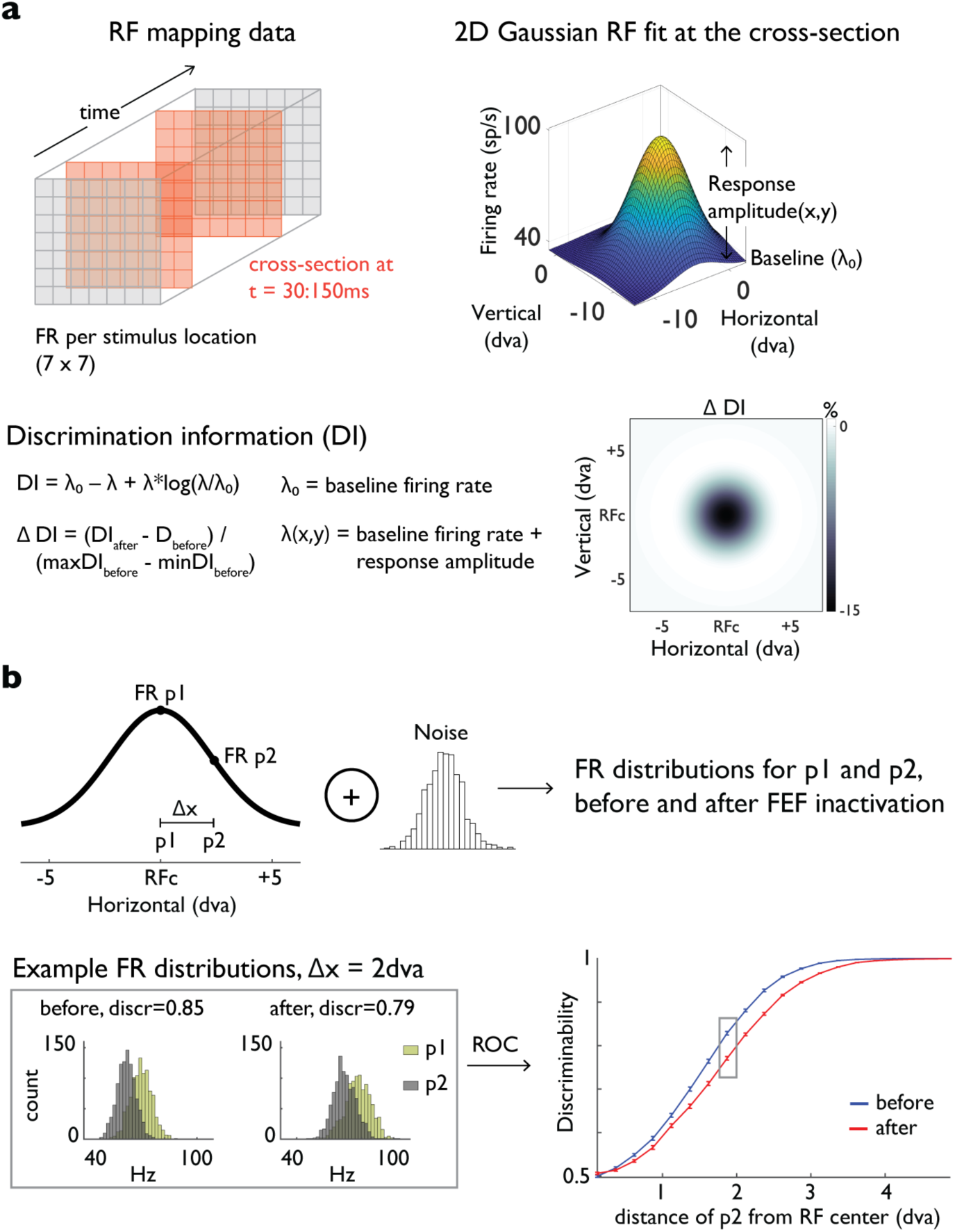
Methods for calculating DI and two-point discriminability. **a**- Top left: Gaussian fits were constructed from responses to probes at each location, 30-150ms after the probe onset. Top right: Illustration of Gaussian RF fit based on the mean of each parameter across the population. Baseline and response amplitude values are indicated. Bottom: DI is calculated using the formula shown, where λ_0_ is baseline, and λ is baseline + response amplitude. Each (x,y) location will have a different λ. DI is calculated from λ_0_ and λ at each location, before and after FEF inactivation. Change in DI is displayed as ΔDI = (DI_before_-DI_after_)/(max DI_before_-min DI_before_). **b**- Top: The population average Gaussian fit was used to generate firing rate distributions for two hypothetical probe locations (p1, p2), before and after FEF inactivation. p1 was always located at the center of the RF and the location of p2 varied. Gaussian noise was added to generate firing rate distributions (n=1000). Bottom left: Example firing rate distributions with p2 at 2dva, before (left) and after FEF inactivation (right). Histograms show number of simulated trials with given firing rates for p1 (green) and p2 (gray). ROC analysis is used to measure an ideal observer’s ability to discriminate between two probe locations based on these firing rate distributions. Bottom right: Discriminability before (blue) and after FEF inactivation (red), as a function of distance of the p2 probe from p1 at the RF center. Box indicates example shown to the left. Error bars show mean ± SE over a bin of probe separations (width = 0.25dva).

### Supplementary Methods

#### Eye calibration

All sessions started with a visual calibration task that allowed us to convert (X, Y) analog signals from the eye tracker to a dva equivalent position on the screen. In this task, the monkey fixated on pseudorandomly presented visual targets for at least 1000ms in order to receive a juice reward. Visual targets were white circles of 1dva diameter on a black background, placed at five different locations on the screen, equally spaced at 10dva from the center of the screen over the horizontal and vertical meridians. The fixation window was ±2 dva. We computed 2D affine transformations to translate the analog (X, Y) eye position to the dva equivalent for further eye movement analyses.

#### V4 RF localization

Prior to data acquisition, we localized retinotopically corresponding V4 and FEF sites. V4 RFs were mapped by assessment of the neuronal firing rate in response to visual stimulus presentation, using a moving white bar stimulus on a black background. In this task, the monkey fixated on a central spot (white circle of 1dva diameter, fixation window ±1.5 dva), while a white bar swept in eight directions over a designated radius centered according to the estimated V4 RF eccentricity. Length and thickness of the bar were adjusted according to the V4 RF eccentricity. The monkey was rewarded with juice if fixation was held throughout the bar sweeping time. V4 responses were sustained to static stimuli, orientation selective, and the RF size increased with RF eccentricity (Gattass et al 1988).

#### FEF RF localization

FEF RFs were localized while exploring the anterior bank of the arcuate sulcus using electrical microstimulation, similar to previous studies (Noudoost et al 2014, Noudoost & Moore 2011a). We delivered biphasic microcurrent pulses (50μA) using a S88 Grass stimulator, via tungsten microelectrodes of 200μm diameter, with epoxylite insulation (FHC, Bowdoin, ME). FEF was identified based on the ability to evoke eye movements of the same direction and amplitude, regardless of the position of the eye on the screen, with currents ≤50μA, while the monkey viewed a black screen with the lights off. We localized FEF sites that when stimulated, elicited eye movements towards the lower visual field, contralaterally, towards the location of the V4 RFs. Landing points of the evoked saccades determined the center of the FEF RF (Fig 1b).

#### Memory guided saccade task to track inactivation effects

In order to track behavioral deficits following FEF inactivation, the animal performed a memory guided saccade task before and after infusion of muscimol in the FEF (Dias & Segraves 1999). In this task, the monkey fixated on a white circle (1dva diameter, 1000ms), and a peripheral visual target (red circle, 1dva diameter, 1000ms) was presented at 1 of 8 possible locations: in the V4 RF, and at 45° intervals in a circle around fixation (Fig 1f and 2a). The monkey maintained fixation while remembering the target location throughout a blank delay period (1000ms + 1:500ms jitter), and after the fixation point disappeared (go cue), executed a saccadic eye movement to the remembered location (target window ±4 dva) to receive a juice reward. Trials were aborted if the animal broke fixation or if blinks were detected.

#### MGS performance and scatter

Behavioral performance on the memory guided saccade task was calculated based on the number of trials in which the saccade was successfully made to the target location, divided by the number of the trials that reached the go cue (or response period) (%). We established a threshold window of 4dva radius around the target location to determine correct trials based on the landing point of the saccade, and implemented an on-line blink detection algorithm that aborted the current trial if any blink was detected at any point from fixation to response period. The scatter of saccade landing points was computed for before and after FEF inactivation trials, based on the saccade endpoints in the response period in the MGS task, for the V4 RF location and 180° opposite to it (Fig S3). Saccade scatter was defined as the average of Euclidean distances between all pairs of saccade endpoints (first saccade after the go cue only, not the corrected one, for all trials that reached the go cue), according to the following equation (Noudoost et al 2014):

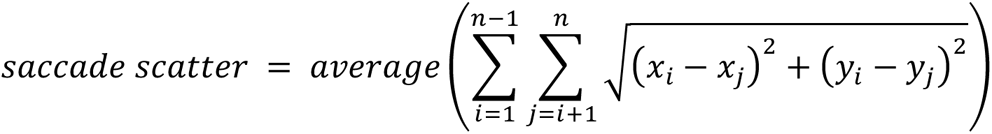

Where the coordinates (x_i_,y_i_) and (x_j_,y_j_) correspond to the eye landing points for a pair of trials i and j.

## References

Acker L, Pino EN, Boyden ES, Desimone R. 2016. FEF inactivation with improved optogenetic methods. Proceedings of the National Academy of Sciences of the United States of America 113: E7297–E306

Adesnik H, Bruns W, Taniguchi H, Huang ZJ, Scanziani M. 2012. A neural circuit for spatial summation in visual cortex. Nature 490: 226–31

Anderson JC, Kennedy H, Martin KAC. 2011. Pathways of Attention: Synaptic Relationships of Frontal Eye Field to V4, Lateral Intraparietal Cortex, and Area 46 in Macaque Monkey. Journal of Neuroscience 31: 10872–81

Armstrong KM, Chang MH, Moore T. 2009. Selection and Maintenance of Spatial Information by Frontal Eye Field Neurons. Journal of Neuroscience 29: 15621–29

Armstrong KM, Fitzgerald JK, Moore T. 2006. Changes in visual receptive fields with microstimulation of frontal cortex. Neuron 50: 791–98

Armstrong KM, Moore T. 2007. Rapid enhancement of visual cortical response discriminability by microstimulation of the frontal eye field. Proceedings of the National Academy of Sciences of the United States of America 104: 9499–504

Balan PF, Gerits A, Zhu Q, Kolster H, Orban GA, et al. 2019. Fast Compensatory Functional Network Changes Caused by Reversible Inactivation of Monkey Parietal Cortex. Cerebral Cortex 29: 2588–606

Bruce CJ, Goldberg ME, Bushnell MC, Stanton GB. 1985. Primate Frontal Eye Fields. II. Physiological and Anatomical Correlates of Electrically Evoked Eye Movements. Journal of Neurophysiology 54: 714–34

Buffalo EA, Fries P, Landman R, Liang H, Desimone R. 2010. A backward progression of attentional effects in the ventral stream. Proceedings of the National Academy of Sciences of the United States of America 107: 361–65

Clark K, Squire RF, Merrikhi Y, Noudoost B. 2015. Visual attention: Linking prefrontal sources to neuronal and behavioral correlates. Progress in neurobiology 132: 59–80

Dias EC, Kiesau M, Segraves MA. 1995. Acute activation and inactivation of macaque frontal eye field with GABA-related drugs. Journal of Neurophysiology 74: 2744–48

Dias EC, Segraves MA. 1999. Muscimol-induced inactivation of monkey frontal eye field: Effects on visually and memory-guided saccades. Journal of Neurophysiology 81: 2191–214

Ekstrom LB, Roelfsema PR, Arsenault JT, Kolster H, Vanduffel W. 2009. Modulation of the Contrast Response Function by Electrical Microstimulation of the Macaque Frontal Eye Field. Journal of Neuroscience 29: 10683–94

Felleman DJ, Van Essen DC. 1991. Distributed Hierarchical Processing in the Primate Cerebral Cortex. Cerebral Cortex 1: 1–47

Gattass R, Sousa APB, Gross CG. 1988. Visuotopic Organization and Extent of V3 and V4 of the Macaque. Journal of Neuroscience 8: 1831–45

Gregoriou GG, Gotts SJ, Zhou HH, Desimone R. 2009. High-Frequency, Long-Range Coupling Between Prefrontal and Visual Cortex During Attention. Science 324: 1207–10

Gregoriou GG, Rossi AF, Ungerleider LG, Desimone R. 2014. Lesions of prefrontal cortex reduce attentional modulation of neuronal responses and synchrony in V4. Nature Neuroscience 17: 1003–11

Hamker FH, Zirnsak M. 2006. V4 receptive field dynamics as predicted by a systems-level model of visual attention using feedback from the frontal eye field. Neural Networks 19: 1371–82

Hwang J, Mitz AR, Murray EA. 2019. NIMH MonkeyLogic: Behavioral control and data acquisition in MATLAB. Journal of Neuroscience Methods 323: 13–21

Lee S-H, Kwan AC, Zhang S, Phoumthipphavong V, Flannery JG, et al. 2012. Activation of specific interneurons improves V1 feature selectivity and visual perception. Nature 488: 379-+

Markov NT, Ercsey-Ravasz M, Lamy C, Gomes ARR, Magrou L, et al. 2013. The role of long-range connections on the specificity of the macaque interareal cortical network. Proceedings of the National Academy of Sciences of the United States of America 110: 5187–92

Markov NT, Vezoli J, Chameau P, Falchier A, Quilodran R, et al. 2014. Anatomy of Hierarchy: Feedforward and Feedback Pathways in Macaque Visual Cortex. Journal of Comparative Neurology 522: 225–59

Merrikhi Y, Clark K, Albarran E, Parsa M, Zirnsak M, et al. 2017. Spatial working memory alters the efficacy of input to visual cortex. Nature Communications 8

Mitchell JF, Sundberg KA, Reynolds JH. 2007. Differential attention-dependent response modulation across cell classes in macaque visual area V4. Neuron 55: 131–41

Monosov IE, Thompson KG. 2009. Frontal Eye Field Activity Enhances Object Identification During Covert Visual Search. Journal of Neurophysiology 102: 3656–72

Moore T, Armstrong KM. 2003. Selective gating of visual signals by microstimulation of frontal cortex. Nature 421: 370–73

Moore T, Fallah M. 2001. Control of eye movements and spatial attention. Proceedings of the National Academy of Sciences of the United States of America 98: 1273–76

Moore T, Tolias AS, Schiller PH. 1998. Visual representations during saccadic eye movements. Proceedings of the National Academy of Sciences of the United States of America 95: 8981–84

Moran J, Desimone R. 1985. Selective attention gates visual processing in the extrastriate cortex. Science 229: 782–84

Mueller A, Krock RM, Shepard S, Moore T. 2020. Dopamine Receptor Expression Among Local and Visual Cortex-Projecting Frontal Eye Field Neurons. Cerebral Cortex 30: 148–64

Nassi JJ, Lomber SG, Born RT. 2013. Corticocortical feedback contributes to surround suppression in V1 of the alert primate. Journal of Neuroscience 33: 8504–U440

Nesse WH, Bahmani Z, Clark K, Noudoost B. 2021. Differential Contributions of Inhibitory Subnetwork to Visual Cortical Modulations Identified via Computational Model of Working Memory. Frontiers in Computational Neuroscience 15

Noudoost B, Clark KL, Moore T. 2014. A Distinct Contribution of the Frontal Eye Field to the Visual Representation of Saccadic Targets. Journal of Neuroscience 34: 3687–98

Noudoost B, Moore T. 2011a. Control of visual cortical signals by prefrontal dopamine. Nature 474: 372–75

Noudoost B, Moore T. 2011b. A reliable microinjectrode system for use in behaving monkeys. Journal of Neuroscience Methods 194: 218–23

Nurminen L, Merlin S, Bijanzadeh M, Federer F, Angelucci A. 2018. Top-down feedback controls spatial summation and response amplitude in primate visual cortex. Nature Communications 9

Pouget P, Emeric EE, Stuphorn V, Reis K, Schall JD. 2005. Chronometry of visual responses in frontal eye field, supplementary eye field, and anterior cingulate cortex. Journal of Neurophysiology 94: 2086–92

Robinson DA, Fuchs AF. 1969. Eye Movements Evoked by Stimulation of Frontal Eye Fields. Journal of Neurophysiology 32: 637-&

Ruff CC, Blankenburg F, Bjoertomt O, Bestmann S, Freeman E, et al. 2006. Concurrent TMS-fMRI and psychophysics reveal frontal influences on human retinotopic visual cortex. Current Biology 16: 1479–88

Schafer RJ, Moore T. 2011. Selective Attention from Voluntary Control of Neurons in Prefrontal Cortex. Science 332: 1568–71

Schall JD. 1991. Neuronal Activity Related to Visually Guided Saccades in the Frontal Eye Fields of Rhesus Monkeys: Comparison with Supplementary Eye Fields. Journal of Neurophysiology 66: 559–79

Schall JD, Morel A, King DJ, Bullier J. 1995. Topography of Visual Cortex Connections with Frontal Eye Field in Macaque: Convergence and Segregation of Processing Streams. Journal of Neuroscience 15: 4464–87

Schmolesky MT, Wang YC, Hanes DP, Thompson KG, Leutgeb S, et al. 1998. Signal timing across the macaque visual system. Journal of Neurophysiology 79: 3272–78

Silvanto J, Lavie N, Walsh V. 2006. Stimulation of the human frontal eye fields modulates sensitivity of extrastriate visual cortex. Journal of Neurophysiology 96: 941–45

Squire RF, Noudoost B, Schafer RJ, Moore T. 2013. Prefrontal Contributions to Visual Selective Attention. Annual Review of Neuroscience*, Vol* 36 36: 451–66

Steinmetz NA, Moore T. 2014. Eye Movement Preparation Modulates Neuronal Responses in Area V4 When Dissociated from Attentional Demands. Neuron 83: 496–506

Taylor PCJ, Nobre AC, Rushworth MFS. 2007. FEF TMS affects visual cortical activity. Cerebral Cortex 17: 391–99

Thompson KG, Bichot NP. 2005. A visual salience map in the primate frontal eye field. Development, Dynamics and Pathology of Neuronal Networks: from Molecules to Functional Circuits 147: 251–62

Ungerleider LG, Galkin TW, Desimone R, Gattass R. 2008. Cortical connections of area V4 in the macaque. Cerebral Cortex 18: 477–99

Vanegas MI, Hubbard KR, Esfandyarpour R, Noudoost B. 2019. Microinjectrode System for Combined Drug Infusion and Electrophysiology. Journal of Visualized Experiments: JoVE

Veniero D, Gross J, Morand S, Duecker F, Sack AT, Thut G. 2021. Top-down control of visual cortex by the frontal eye fields through oscillatory realignment. Nature communications 12: 1757–57

Vinck M, Womelsdorf T, Buffalo EA, Desimone R, Fries P. 2013. Attentional Modulation of Cell-Class-Specific Gamma-Band Synchronization in Awake Monkey Area V4. Neuron 80: 1077–89

Wardak C, Ben Hamed S, Olivier E, Duhamel J-R. 2012. Differential effects of parietal and frontal inactivations on reaction times distributions in a visual search task. Frontiers in integrative neuroscience 6: 39–39

Wardak C, Ibos G, Duhamel JR, Olivier E. 2006. Contribution of the monkey frontal eye field to covert visual attention. Journal of Neuroscience 26: 4228–35

Williams GV, Goldman-Rakic PS. 1995. Modulation of memory fields by dopamine D1 receptors in prefrontal cortex. Nature 376: 572–75

Zanto TP, Rubens MT, Thangavel A, Gazzaley A. 2011. Causal role of the prefrontal cortex in top-down modulation of visual processing and working memory. Nature Neuroscience 14: 656–U156

